# Cellular sialoglycans are differentially required for endosomal and cell-surface entry of SARS-CoV-2

**DOI:** 10.1101/2024.06.24.600376

**Authors:** Kimberley C. Siwak, Emmanuelle V. LeBlanc, Heidi M. Scott, Youjin Kim, Isabella Pellizzari-Delano, Alice M. Ball, Nigel J. Temperton, Chantelle J. Capicciotti, Che C. Colpitts

**Affiliations:** Department of Biomedical and Molecular Sciences, Queen’s University, Kingston, ON K7L 3N6; Viral Pseudotype Unit, Medway School of Pharmacy, University of Kent and Greenwich at Medway, Chatham, UK; Department of Chemistry, Queen’s University, Kingston, ON K7L 3N6; Department of Surgery, Queen’s University, Kingston, ON K7L 3N6

## Abstract

Cell entry of severe acute respiratory coronavirus-2 (SARS-CoV-2) and other CoVs can occur via two distinct routes. Following receptor binding by the spike glycoprotein, membrane fusion can be triggered by spike cleavage either at the cell surface in a transmembrane serine protease 2 (TMPRSS2)-dependent manner or within endosomes in a cathepsin-dependent manner. Cellular sialoglycans have been proposed to aid in CoV attachment and entry, although their functional contributions to each entry pathway are unknown. In this study, we used genetic and enzymatic approaches to deplete sialic acid from cell surfaces and compared the requirement for sialoglycans during endosomal and cell-surface CoV entry, primarily using lentiviral particles pseudotyped with the spike proteins of different sarbecoviruses. We show that entry of SARS-CoV-1, WIV1-CoV and WIV16-CoV, like the SARS-CoV-2 omicron variant, depends on endosomal cathepsins and requires cellular sialoglycans for entry. Ancestral SARS-CoV-2 and the delta variant can use either pathway for entry, but only require sialic acid for endosomal entry in cells lacking TMPRSS2. Binding of SARS-CoV-2 spike protein to cells did not require sialic acid, nor was sialic acid required for SARS-CoV-2 entry in TMRPSS2-expressing cells. These findings suggest that cellular sialoglycans are not strictly required for SARS-CoV-2 attachment, receptor binding or fusion, but rather promote endocytic entry of SARS-CoV-2 and related sarbecoviruses. In contrast, the requirement for sialic acid during entry of MERS-CoV pseudoparticles and authentic HCoV-OC43 was not affected by TMPRSS2 expression, consistent with a described role for sialic acid in merbecovirus and embecovirus cell attachment. Overall, these findings clarify the role of sialoglycans in SARS-CoV-2 entry and suggest that cellular sialoglycans mediate endosomal, but not cell-surface, SARS-CoV-2 entry. Thus, it may be important to consider both cell entry pathways when developing sarbecovirus entry inhibitors targeting virus-sialoglycan interactions.

**Author summary:** The COVID-19 pandemic, caused by SARS-CoV-2, has resulted in over 676 million infections and 6.8 million deaths so far, demonstrating the threat posed by emerging CoVs. In humans, SARS-CoV-2 and related coronaviruses cause respiratory tract infections, such as the common cold, as well as more severe disease in some individuals. To prepare for future outbreaks, conserved steps in the CoV replication could be considered for antiviral prophylactic or therapeutic approaches. One such process is CoV cell entry, which occurs via two main routes: At the cell surface or within endosomes. Cellular receptors, proteases and complex sugars, known as glycans, mediate CoV entry steps. In this study, we compared the role of a specific glycan subset, sialoglycans, in endosomal and cell surface CoV entry. We show that sialoglycans are required for entry of various CoVs that are mainly dependent on the endosomal route, but in the case of SARS-CoV-2, sialoglycans were not required when the cell-surface entry route was available. Our findings contribute to understanding the mechanisms of CoV entry, which could inform development of pan-CoV antivirals that target CoV entry steps.

## Introduction

The COVID-19 pandemic highlights our vulnerability to emerging coronaviruses (CoVs). With three outbreaks of highly pathogenic CoVs in the past 20 years, there is increasing risk of another cross-species transmission event from zoonotic reservoirs, such as bats (1), into human populations. In particular, bats harbour many novel SARS-like and MERS-like CoVs, which could pose a threat for spillover to humans (1–3), including WIV1-CoV and WIV16-CoV, which are closely related to SARS-CoV-1 and are suggested to be poised for human emergence (4, 5). Novel antiviral countermeasures that are broadly protective against CoVs could help protect against future pandemic CoV threats that may emerge.

CoV cell entry, a critical determinant of viral tropism, pathogenesis, and cross-species transmission, could serve as a target for potential broadly acting antiviral therapies. The spike (S) protein mediates entry into cells by interacting with specific host cell entry receptors. Many CoVs, such as SARS-CoV-2 and SARS-CoV-1, as well as the pre-emergent bat CoVs, WIV1-CoV and WIV16-CoV, have evolved to use human angiotensin-converting enzyme 2 (ACE2) as a host receptor (6–11), while the MERS-CoV receptor is dipeptidyl peptidase 4 (DPP4) (12). In addition to receptor binding, proteolytic cleavage of the S protein is required for fusion and entry (13). Several proteases can facilitate these proteolytic cleavage events, which determines the cell entry route. Proteolytic cleavage can be mediated by the conserved cell surface transmembrane protease serine 2 (TMPRSS2) (14, 15), which is found at the plasma membrane and therefore its cleavage of the CoV spike protein promotes fusion and entry at the cell surface. Cathepsin L, a non-specific cysteine protease, has also been implicated in CoV entry (16–19). Cathepsin L is found in endo/lysosomal compartments, requiring virions to be endocytosed prior to cathepsin L-mediated cleavage (16–20). This alternate entry route is termed the endosomal entry pathway (15–17, 21).

The spike proteins of some CoVs, such as SARS-CoV-2 and MERS-CoV, have a multibasic furin cleavage site at the S1/S2 junction, enabling priming of the entry process through cleavage by furin, a specialized serine endoprotease (22). Furin pre-activates the S protein during egress, priming the S protein for the second required cleavage event at S2′ to expose the fusion peptide (23). Furin is thought to act in conjunction with TMPRSS2-mediated S2′ cleavage, promoting cell surface entry over the endosomal pathway (24). For example, early SARS-CoV-2 isolates, which contain a furin cleavage site at spike S1/S2, favour TMPRSS2-mediated S2′ cleavage for cell entry, while SARS-CoV-1, which lacks a spike furin cleavage site, is more dependent on cathepsin L (18, 19, 25).

The S protein also interacts with cell surface glycans to facilitate initial interactions with the host cell (14, 26). Sialic acid, a terminal glycan epitope found abundantly on cellular complex glycans, has been described as a potential co-receptor facilitating attachment of several human CoVs, including SARS-CoV-2, MERS-CoV, HCoV-OC43 and HCoV-HKU1 (27–32). However, while binding of sialic acid by these CoV S proteins has been well-documented through in vitro assays (33–36), the functional role of sialic acid during entry of SARS-CoV-2 has remained less clear. Given previous literature suggesting that sialic acid contributes to internalization of other viruses (37, 38), we sought to characterize the role of cellular sialic acid in the different entry routes used by CoVs. Here, we evaluated the functional roles of sialoglycans during entry of recently emerged beta-CoVs, including SARS-CoV-1, SARS-CoV-2 variants and MERS-CoV, as well as pre-emergent bat CoVs (WIV1-CoV and WIV16-CoV), using lentiviral pseudoparticles as a surrogate model for entry. Interestingly, the requirement for sialic acid during CoV entry is at least partially dependent on the entry route. We show that endosomal entry of bat CoVs WIV1-CoV and WIV16-CoV, as well as SARS-CoV-1 and SARS-CoV-2 omicron variant, confers increased dependence on sialic acid during entry. In contrast, the requirement for sialic acid for entry of SARS-CoV-2 Hu-1 and delta variant varies depending on the cell type and relative abundance of specific proteases, with TMPRSS2 expression abrogating the requirement for sialic acid during entry. These findings suggest a role for sialic acid in mediating endosomal, cathepsin-dependent sarbecovirus entry, but not cell surface, TMPRSS2-mediated entry. On the other hand, entry of MERS-CoV pseudoparticles and authentic HCoV-OC43 relied on sialic acid regardless of TMPRSS2 expression, consistent with a role for sialic acid in attachment of merbecoviruses (33) and embecoviruses (39). Overall, this study provides new insight into the roles of cellular sialoglycans during sarbecovirus entry and contributes to understanding the entry processes of recently emerged and pre-emergent CoVs.

## Results

### Infectivity of pre-emergent bat CoVs and recently emerged CoVs in A549-derived and Calu-3 cell lines

We used lentiviral pseudoparticles (40–42) as a surrogate model to assess and compare the cell entry routes of highly pathogenic human CoVs and bat CoVs. Previous literature has demonstrated the ability of CoVs to infect a variety of human cell lines, including the alveolar epithelial A549 cell line engineered to express appropriate receptors (ACE2 or DPP4) and the Calu-3 lung epithelial cell line, which naturally expresses ACE2, DPP4 and TMPRSS2 (35, 43–47). We generated A549 cell lines that stably express ACE2 (A549-A) or DPP4 (A549-D) in the presence or absence of TMPRSS2. We confirmed expression of ACE2 or DPP4 by western blot **(Fig 1A)** and TMPRSS2 by flow cytometry **(Fig 1B)**. We first compared MERS-CoV pseudoparticle entry in A549-D, A549-DPP4-TMPRSS2 (A549-DT) and Calu-3 cells and noted similar infectivity across all cell lines **(Fig 1C)** when normalized to the infectivity of naked pseudoparticles (no env; lacking envelope protein) in each cell line. Similarly, we tested susceptibility of A549-A, A549-ACE2-TMPRSS2 (A549-AT) and Calu-3 cells to lentiviral particles pseudotyped with the spike proteins of ancestral SARS-CoV-2 (Hu-1), B.1.617.2 (delta) and B.1.1.529 (omicron) variants, or SARS-CoV-1 and related bat CoVs WIV1-CoV and WIV16-CoV **(Fig 1D)**. We confirmed the ACE2 dependence of WIV1-CoV and WIV16-CoV **(S1 Fig)**. While all tested SARS-like CoV pseudoparticles robustly infected A549-A and A549-AT cells, infectivity of SARS-CoV-2 omicron, SARS-CoV-1, WIV1 and WIV16 was profoundly reduced in Calu-3 cells **(Fig 1D)**. Since Calu-3 cells express higher levels of TMPRSS2, but relatively low levels of cathepsin L (15), our findings are consistent with a role for TMPRSS2 in aiding entry of some SARS-CoV-2 variants, but not SARS-CoV-1 or bat CoVs that lack a furin cleavage site and are thus likely more dependent on endosomal entry. TMPRSS2 expression in A549 cells enhanced entry of SARS-CoV-2 Hu1 and delta, but not SARS-CoV-2 omicron or SARS1-like CoVs **(Fig 1D)**. Importantly, infectivity of control VSV-G-pseudotyped lentiviral particles was similar in A549-A and A549-AT cells **(Fig 1D)**, confirming that differences we observed are not the result of downstream steps of lentiviral transduction. Infectivity of VSV-G pseudoparticles was enhanced in Calu-3 cells compared to A549-A and A549-AT cells **(Fig 1D)**, indicating that the impaired entry of some CoV pseudoparticles in Calu-3 cells is not the result of intrinsic resistance of these cells to lentiviral transduction.

**Figure 1.**
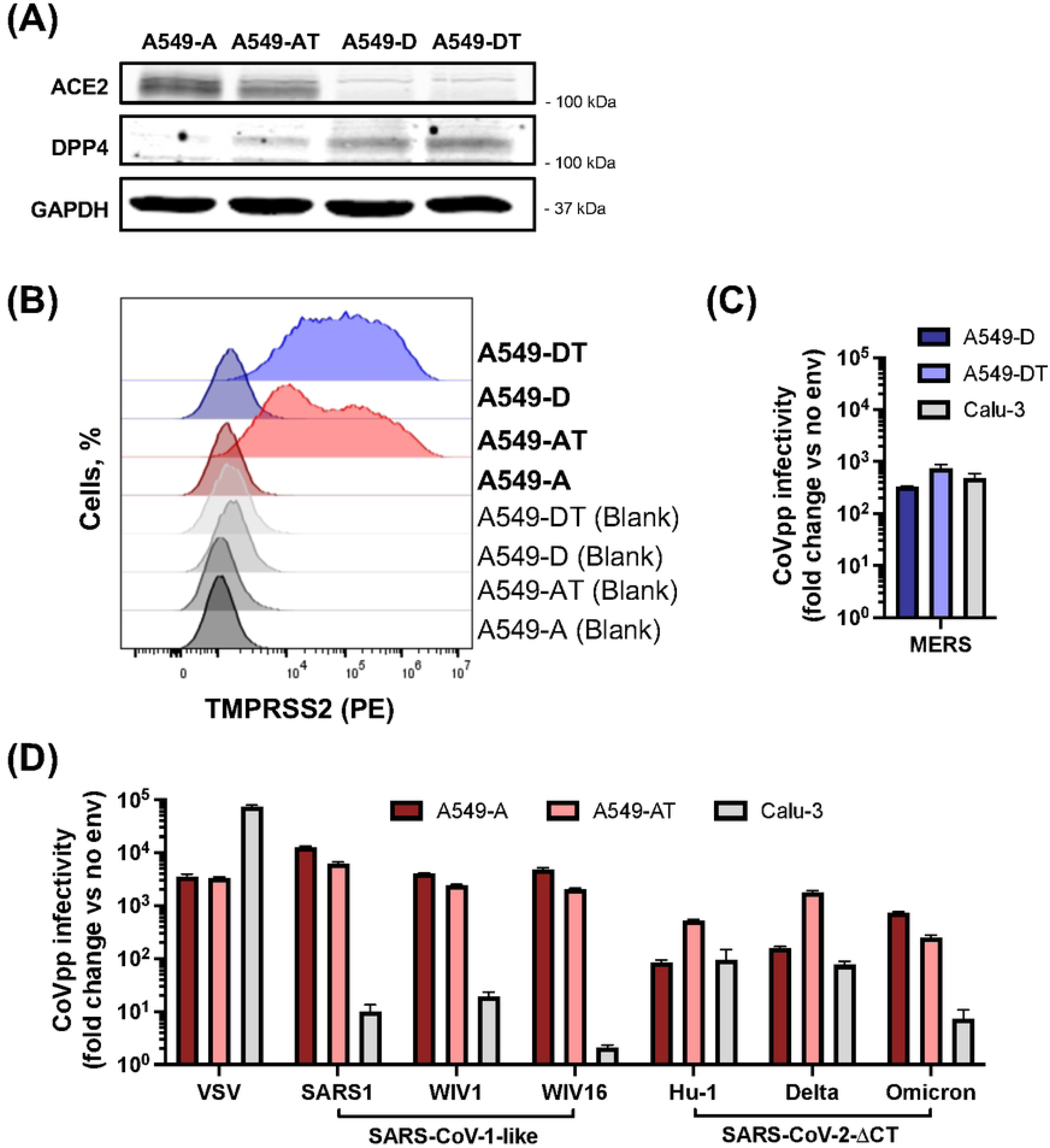
Infectivity of CoVpp in A549-derived and Calu-3 cell lines. **(A-B)** A549 cells were transduced to stably overexpress ACE2 (A549-A), DPP4 (A549-D) in the absence or presence of TMPRSS2 (A549-AT and A549-DT). ACE2 and DPP4 expression was confirmed by western blot **(A)**, while TMPRSS2 expression was confirmed using flow cytometry **(B)**. **(C-D)** A549-derived or Calu-3 cells were inoculated with lentiviral particles pseudotyped with the spike proteins of MERS-CoV **(C)**, VSV, SARS-CoV-1, WIV1-CoV, WIV16-CoV, SARS-CoV-2 Hu1, SARS-CoV-2 delta or SARS-CoV-2 omicron **(D)** or pseudoparticles lacking envelope protein (no env) **(C-D)** for 2 h, then incubated for an additional 72 h, at which point luciferase activity was measured to assess pseudoparticle entry. The data are expressed as fold change relative to the luciferase signal obtained with no envelope. Graphs show mean +/-SEM from three independent experiments performed in triplicate.

We compared entry of SARS-CoV-2 Hu-1 pseudoparticles with a 19-amino acid C-terminally truncated spike protein (SARS-CoV-2-ΔCT) and full-length spike protein (SARS-CoV-2-FL). Consistent with previous literature (40, 48–50), we noted that SARS-CoV-2-FL pseudoparticles had poorer infectivity in A549-A cells, but enhanced infectivity in Calu-3 cells relative to the SARS-CoV-2-ΔCT particles **(S2 Fig)**. While SARS-CoV-2-ΔCT spike proteins are commonly utilized to improve lentiviral vector infectivity (40, 48–50), TMPRSS2-expressing Calu-3 cells appear to be more susceptible to the SARS-CoV-2-FL spike pseudoparticles, as the C-terminal truncation may result in less furin processing and therefore enhanced cathepsin L dependence, rather than TMPRSS2 dependence, during viral entry (44, 51, 52). Therefore, for SARS-CoV-2 Hu1 experiments, we used FL spike pseudoparticles for subsequent entry assays in Calu-3 cells, while ΔCT spike pseudoparticles were used for assays in A549-derived cells.

### Entry route preference of SARS-CoV-2 and MERS-CoV depends on TMPRSS2 expression

We next evaluated the entry route preferences of CoVpp in our cell models using camostat mesylate or E64d to inhibit TMPRSS2 (cell surface entry) or cathepsin L (endosomal entry), respectively **(Fig 2A)** (53, 54). A549-derived cells were pre-treated with DMSO vehicle, camostat (25 μM) or E64d (10 μM) for 1 hour at 37°C prior to inoculation with CoVpp. For MERS-CoVpp, entry was inhibited by E64d, but not camostat, treatment in A549-D cells, but the opposite was observed in A549-DT and Calu-3 cells, confirming that MERS-CoV entry proceeds through the endosomal pathway in A549-D cells, but through the cell-surface TMPRSS2-mediated pathway in TMPRSS2-expressing A549-DT and Calu-3 cells **(Fig 2B)**. For SARS-CoV-2, entry of Hu1, delta and omicron pseudoparticles in A549-A cells was inhibited by treatment with E64d, but not camostat **(Fig 2C)**, consistent with the lack of TMPRSS2 expression in these cells **(Fig 1B)**. In A549-AT cells, on the other hand, entry of SARS-CoV-2 Hu-1 and delta pseudoparticles was strongly reduced by camostat, but to a much lesser extent by E64d **(Fig 2C)**, indicating a preference for cell-surface TMPRSS2-mediated entry when TMPRSS2 is present. Entry of SARS-CoV-2 omicron pseudoparticles in A549-AT cells was partially inhibited by both E64d or camostat, but more potently inhibited by E64d, indicating that the endosomal pathway is preferred even in the presence of TMPRSS2 **(Fig 1C)**, which is consistent with prior literature (54). Finally, entry of SARS-CoV-1, WIV1-CoV and WIV16-CoV spike-pseudotyped particles was inhibited by E64d, whereas camostat did not significantly affect entry of SARS-CoV-1 or WIV16-CoV pseudoparticles, although did weakly inhibit entry of WIV1-CoV pseudoparticles **(Fig 2D)**. Overall, we concluded that entry of SARS-CoV-1, WIV1-CoV and WIV16-CoV occurs predominantly through the endosomal route even in the presence of TMPRSS2.

**Figure 2.**
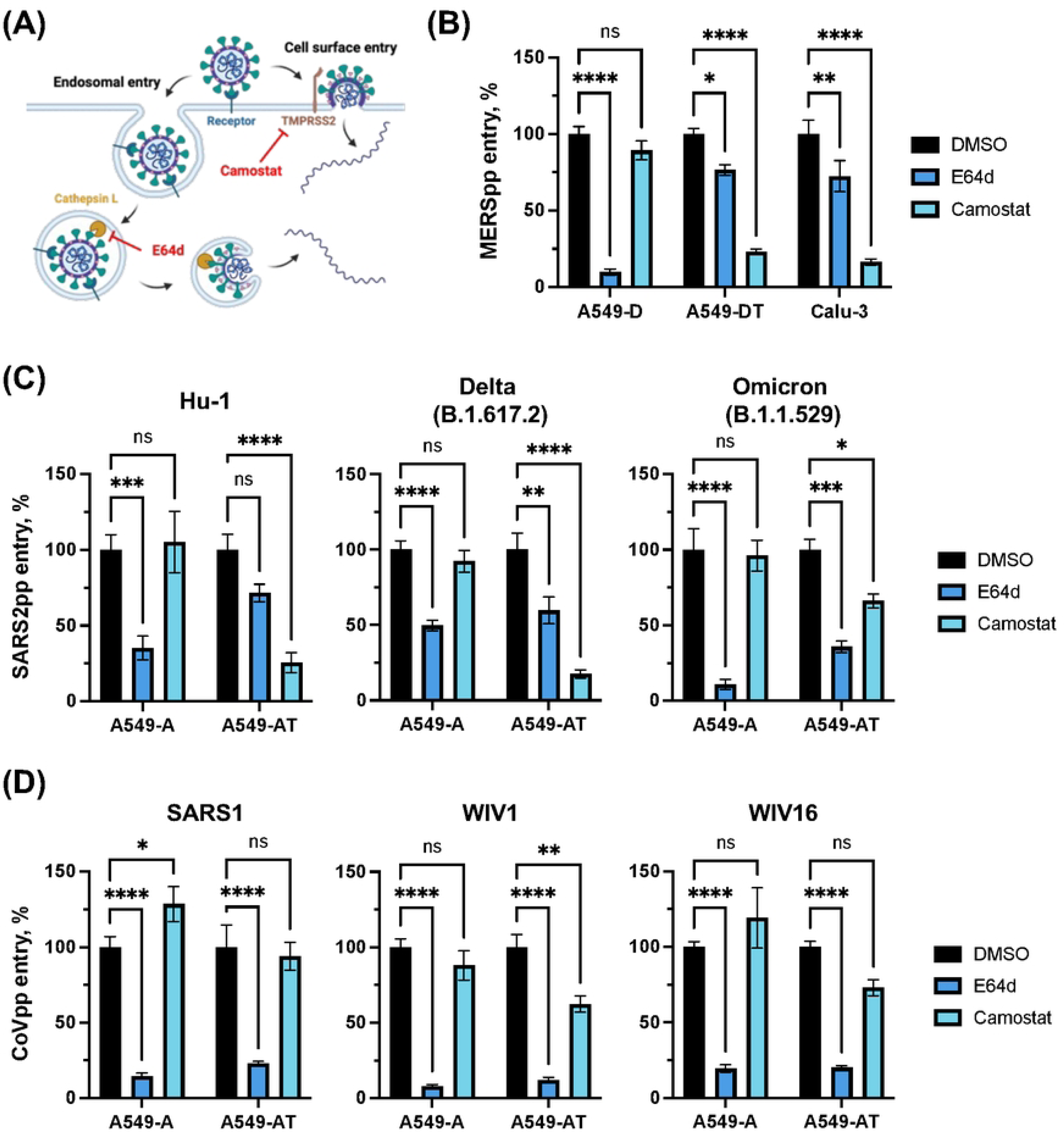
The entry route of SARS-CoV-2 and MERS-CoV depends on TMPRSS2 expression in A549 cells. **(A)** Schematic showing cell-surface TMPRSS2-mediated CoV entry compared to endosomal cathepsin-mediated entry. Camostat mesylate inhibits the TMPRSS2 pathway, while E64d inhibits the endosomal pathway. **(B-D)** A549-D, A549-DT or Calu-3 cells **(B)** or A549-A or A549-AT cells **(C-D)** were pre-treated for 1 h at 37°C with DMSO, camostat (25 μM) or E64d (10 μM) diluted in media to the indicated concentrations, then infected with MERS-CoVpp **(B)**, SARS-CoV-2pp **(C)** or SARS-CoV-1-like pseudoparticles **(D)** for 2 h at 37°C. Inocula were removed and cells were incubated in complete media for 72 h, at which point luciferase activity was measured to assess viral entry. Data are expressed as percentage relative to DMSO control. Graphs show mean +/- SEM from at least three independent experiments performed in triplicate. Statistical significance was assessed by two-way ANOVA (*p<0.05; **p<0.01; ***p<0.001; ****p<0.0001; ns, not significant).

### Sialic acid differentially contributes to sarbecovirus entry depending on entry route

We first used genetic and enzymatic approaches to evaluate the role of sialic acid during CoV entry in A549-A cells, where the endosomal entry route is predominant **(Fig 2C-D)**. We used CRISPR/Cas9 to generate cells lacking cytidine monophosphate N-acetylneuraminic acid synthetase (CMAS), an enzyme that catalyzes the conversion of N-acetylneuraminic acid (Neu5Ac) to cytidine 5’-monophosphate N-acetylneuraminic acid (CMP-Neu5Ac), the essential nucleotide sugar donor required for the synthesis of sialylated glycans (55). Alternatively, we used the broad acting sialidase NanH from *Clostridium perfringens* to enzymatically remove terminal sialic acid epitopes from cell surfaces (56). Cells were pre-treated for 30 minutes at 37°C with NanH diluted to 50 μg/mL in serum-free F12-K media. NanH was then removed, and cells were washed with PBS. We confirmed absence of sialic acid in CMAS knockout (KO) A549-A cells, or in WT A549-A cells treated with NanH, by lectin staining and flow cytometry using fluorescently labelled sialic acid-binding lectin *Sambucus nigra* agglutinin (SNA-FITC) and terminal-galactose-binding *Erythrina cristagalli* lectin (ECL-FITC) **(Fig 3A)**. SNA-FITC binding to cells was decreased in the absence of sialic acid, with a concomitant increase in ECL-FITC binding reflecting increased exposure of galactose in the absence of terminal sialic acid **(Fig 3A)**. Residual binding signal after NanH treatment or in CMAS KO cell line may be attributed to non-specific background binding which has been observed in previous studies (57, 58). We tested the role of sialic acid for spike glycoprotein (SGP) binding to these cells using flow cytometry. A549-A WT or CMAS KO cells were treated with NanH or mock-treated and incubated with soluble recombinant SARS-CoV-2 Hu-1 SGP. SGP bound A549-ACE2 cells similarly regardless of NanH treatment or CMAS KO **(Fig 3B)**, indicating that terminal sialic acid is not required for SARS-CoV-2 spike binding to A549-A cells. As a control, treatment with soluble heparin did inhibit spike binding to cells **(Fig 3C)**, reflecting the established role of heparan sulfate proteoglycans (HSPGs) in mediating SARS-CoV-2 attachment (26).

**Figure 3.**
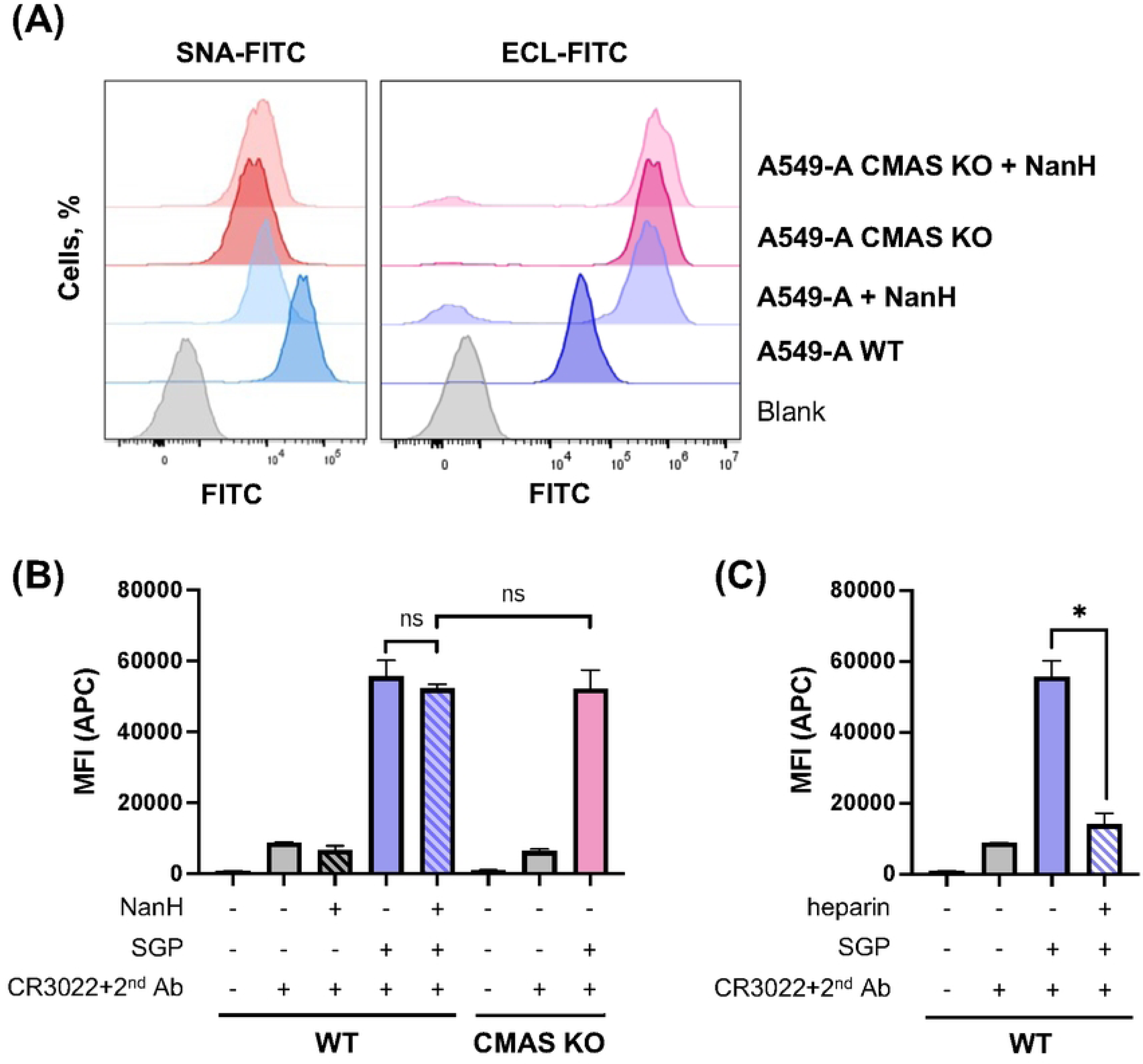
Cell surface sialic acid is not required for SARS-CoV-2 spike glycoprotein binding. **(A)** The presence of cell surface sialic acid was confirmed by flow cytometry using *Sambucus nigra* agglutinin (SNA-FITC) and *Erythrina cristagalli* lectin (ECL-FITC). SNA binds to sialylated glycans, and a decrease is observed upon loss of sialic acid; ECL binds to terminal galactosylated glycans, and an increase is observed upon loss of sialic acid. **(B-C)** Soluble recombinant spike glycoprotein (SGP) binding to A549-A and A549-A CMAS KO cells assessed by flow cytometry. Enzymatic desialylation was achieved by incubating cells with NanH for 30 min. Heparin inhibition was performed by pre-incubating SGP with heparin 10 min at 37 °C prior to the addition to cells. Graphs show mean +/- SEM from an experiment performed in duplicate. Statistical significance was assessed by Student’s t-test (*p<0.05; **p<0.01; ns, not significant).

To determine if the loss of terminal sialic acid epitopes affected SARS-CoV-2 and other CoV entry at a post-binding step, we performed pseudoparticle entry assays in CMAS KO A549-A cells compared to WT A549-A cells. Entry of all sarbecovirus pseudoparticles was significantly reduced in CMAS KO cells compared to parental A549-A cells **(Fig 4A)**. We also observed that entry of VSV G-pseudotyped lentiviral particles was decreased in the CMAS KO cells by approximately 50%, which could support a proposed role for sialic acid in VSV entry (59) Interestingly, we noted differences in the inhibitory effect of CMAS KO for different CoV pseudoparticles, with entry of SARS1-like CoVs as well as the SARS-CoV-2 omicron variant being more profoundly affected by the absence of sialic acid (>90% inhibition) than SARS-CoV-2 Hu-1 or delta strains (∼50% inhibition) **(Fig 4A)**, which reflected the inhibition pattern when endosomal entry was blocked via E64d treatment **(Fig 2A-B)**. Therefore, we directly compared the effect of E64d on entry of SARS-CoV-2 pseudoparticles in WT and CMAS KO A549-A cells. E64d treatment in WT cells reduced entry of SARS-CoV-2 Hu-1, delta and omicron to a similar extent as in DMSO-treated CMAS KO cells **(Fig 4B)**. Interestingly, while SARS-CoV-2 entry was already inhibited in CMAS KO cells, there was no further inhibitory effect of E64d in CMAS KO cells **(Fig 4B)**, which could suggest that the absence of sialic acid and inhibition of cathepsins may affect the same step.

**Figure 4.**
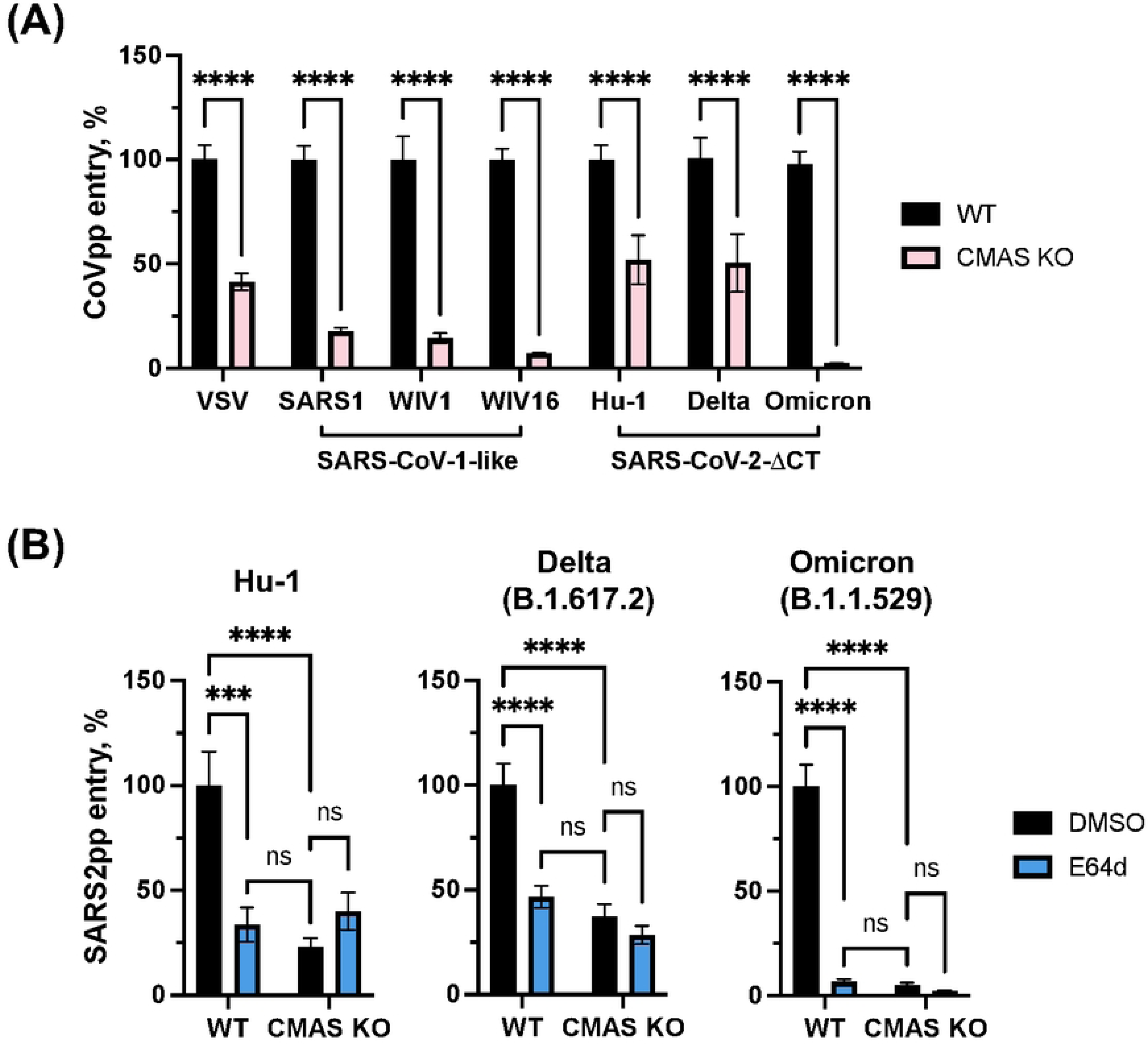
Sarbecovirus entry is reduced in cells lacking terminal sialic acid. **(A)** A549-A WT or CMAS KO cells were inoculated with the indicated pseudoparticles for 2 h at 37°, at which point the inocula were removed and cells were overlaid with complete media. After 72 h, lysates were collected to measure luciferase reporter activity to assess pseudoparticle entry. **(B)** A549-A WT or CMAS KO cells were pre-treated with DMSO vehicle or E64d (10 μM) for 1 h at 37°C, then inoculated with pseudoparticles as described above. Data are expressed as percentage relative to WT cells **(A)** or DMSO-treated control cells **(B)**. **(A-B)** Graphs show mean +/- SEM from at least three independent experiments performed in triplicate. Statistical significance was assessed by two-way ANOVA (***p<0.001; ****p<0.0001; ns, not significant).

### TMPRSS2 expression decreases sialic acid dependence during SARS-CoV-2, but not MERS-CoV or HCoV-OC43, entry

Since SARS-CoV-1-like CoVs and the SARS-CoV-2 omicron variant predominantly use the endosomal entry route **(Fig 2A-B)** and were more profoundly affected by the absence of sialic acid than SARS-CoV-2 Hu-1 or delta **(Fig 3C)**, we hypothesized that sialic acid contributes more to endosomal cathepsin-mediated entry than cell-surface TMPRSS2-mediated entry. To test this hypothesis, we compared the effect of NanH treatment in cells that express or lack TMPRSS2, focusing on SARS-CoV-2 Hu-1, delta and MERS-CoV due to their flexibility in entry route depending on TMPRSS2 expression **(Fig 2B-C)**. We first confirmed that 30-minute pre-treatment with 50 μg/mL NanH effectively removes sialic acid on Calu-3 cells by staining with sialic acid-binding SNA-FITC and terminal-galactose-binding ECL-FITC lectins. As expected, immunofluorescence microscopy confirmed the removal of sialic acid from A549 and Calu-3 cells **(Fig 5A, S3 Fig)**.

**Figure 5.**
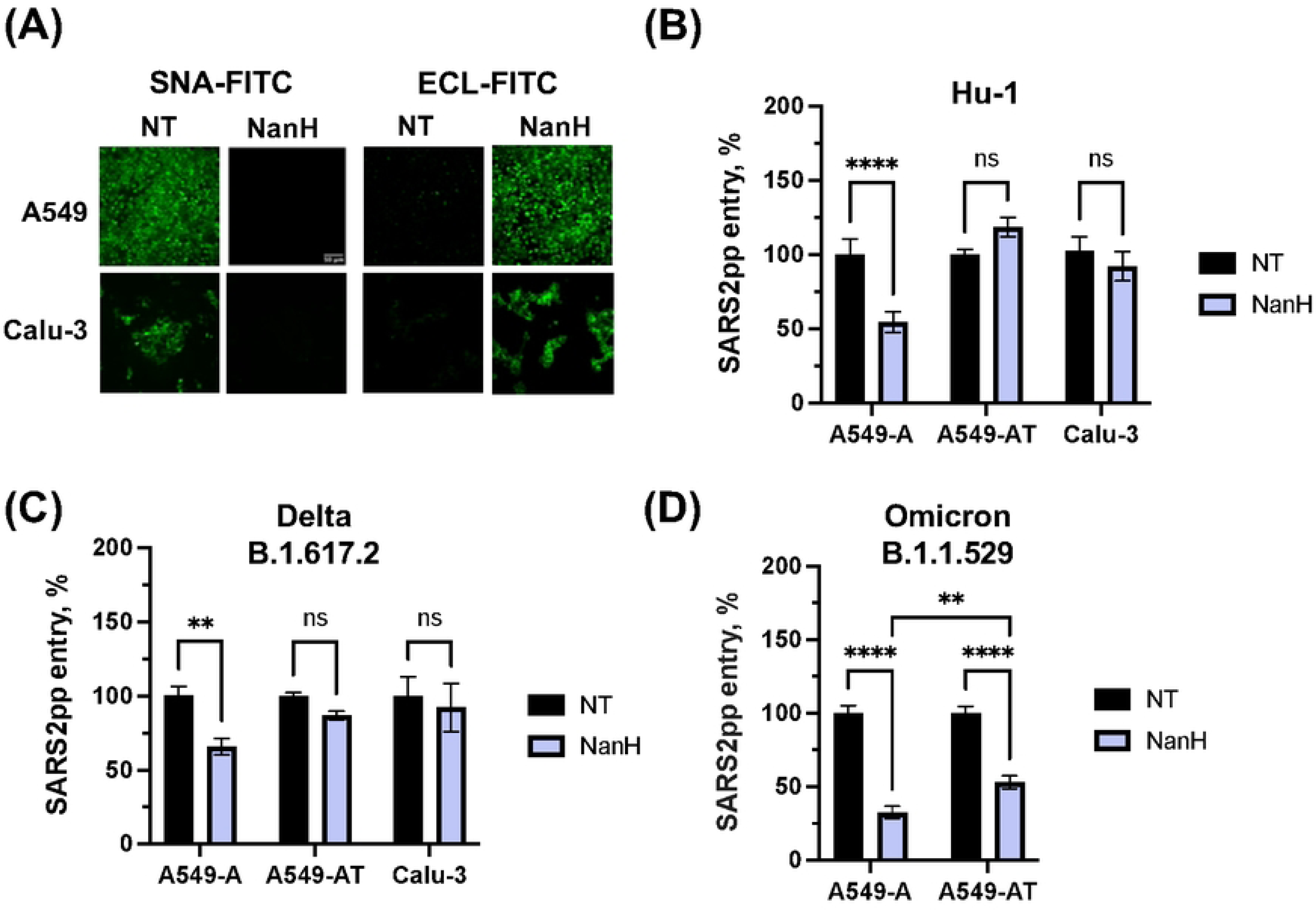
Sialic acid is not required for entry of SARS-CoV-2 Hu1 and delta variant in cells expressing TMPRSS2. **(A-D)** A549-derived cells or Calu-3 cells were pre-treated with NanH diluted to 50 μg/mL in serum-free media for 30 minutes at 37°C. Cells were then washed and processed for fluorescence microscopy **(A)** or inoculated with SARS-CoV-2 pseudoparticles **(B-D)**. **(A)** NanH-treated cells were stained with SNA-FITC (binds sialic acid) or ECL-FITC (binds galactose) diluted to final concentration of 20 μg/mL in PBS, then washed with PBS and imaged by fluorescence microscopy (10X magnification; scale bar, 50 μm). Lectin staining confirmed removal of sialic acid by the NanH treatment. **(B-D)** NanH-treated cells were inoculated with the indicated SARS-CoV-2 pseudoparticles for 2 h at 37°C, then washed and incubated in complete media for 72 h. Luciferase activity was then measured to assess viral entry. Data are expressed as percentage relative to DMSO control. **(B-D)** Graphs show mean +/- SEM from at least three independent experiments performed in triplicate. Statistical significance was assessed by two-way ANOVA (**p<0.01; ****p<0.0001; ns, not significant).

We then performed CoVpp entry assays to determine if the loss of terminal sialic acid epitopes differentially affected SARS-CoV-2 entry in TMPRSS2-lacking A549-A cells (where the endosomal entry route is preferred) compared to TMPRSS2-expressing A549-AT and Calu-3 cells (where the cell surface entry route is preferred). We focused on SARS-CoV-2 due to its flexibility in entry route depending on TMPRSS2 expression **(Fig 2C)**. Reflecting our observation in CMAS KO A549-A cells **(Fig 4A)**, removal of sialic acid by NanH pre-treatment decreased entry of SARS-CoV-2 Hu-1 and delta pseudoparticles in A549-A cells by approximately 50% **(Fig 5B-C)**. However, NanH pre-treatment did not inhibit entry of SARS-CoV-2 Hu-1 and delta pseudoparticles in TMPRSS2-expressing A549-AT or Calu-3 cells **(Fig 5B-C)**, indicating that use of TMPRSS2 and the cell-surface entry route enables entry in the absence of sialic acid. Interestingly, NanH pre-treatment similarly inhibited SARS-CoV-2 omicron entry in A549-A and A549-AT cells, although was slightly less inhibitory in A549-AT cells **(Fig 5D)**, perhaps reflecting the increased dependence of SARS-CoV-2 omicron on the endosomal entry route compared to Hu-1 and delta variants **(Fig 2B)**. Due to very low infectivity of SARS-CoV-2 omicron pseudoparticles in Calu-3 cells **(Fig 1D)**, we were unable to reliably infect Calu-3 cells with omicron pseudoparticles and thus could not assess the effect of NanH on omicron entry in Calu-3 cells.

Finally, we assessed the role of sialic acid in MERS-CoV entry in A549-D cells (endosomal entry) compared to TMPRSS2-expressing A549-DT and Calu-3 cells (cell surface entry). Interestingly, MERS-CoV entry was reduced by NanH pre-treatment in both A549-D and A549-DT cells, but not in Calu-3 cells **(Fig 6A)**. Given the discrepant results in A549-DT and Calu-3 cells, which both support cell-surface entry **(Fig 2B)**, we sought to functionally confirm the absence of sialic acid following NanH treatment of Calu-3 cells by infecting cells with influenza A virus (IAV; strain A/New York/18/2009; H1N1), which uses α2,6-linked sialic acid as a receptor (60). NanH-treated cells were infected with 3×10^3^, 3×10^4^, or 3×10^5^ plaque-forming units (pfu) of IAV, corresponding to an approximate multiplicity of infection (MOI) of 0.01, 0.1 or 1 pfu/cell. At 8 hours post-infection, cells were lysed to measure IAV *M* gene expression by RT-qPCR. As expected, a reduction in IAV infection was observed following NanH pre-treatment at all virus doses **(S3 Fig)**, confirming the functional absence of sialic acid following NanH treatment. We noted that the absence of sialic acid was less inhibitory when increasing amounts of virus were added, although infection was still inhibited even at an MOI of 1. Using a transfer plasmid encoding ZsGreen in our pseudoparticle assays, we established that between 1 and 10% of cells were transduced with MERS-CoVpp (data not shown), reflecting the lower MOI conditions where IAV infection was robustly inhibited in NanH-treated cells. However, despite effective removal of sialic acid from Calu-3 cells as measured by lectin staining **(Fig 5A)** and IAV infection **(S3 Fig)**, sialic acid did not appear to play a role in MERS-CoV pseudoparticle entry in Calu-3 cells **(Fig 6A)**.

**Figure 6.**
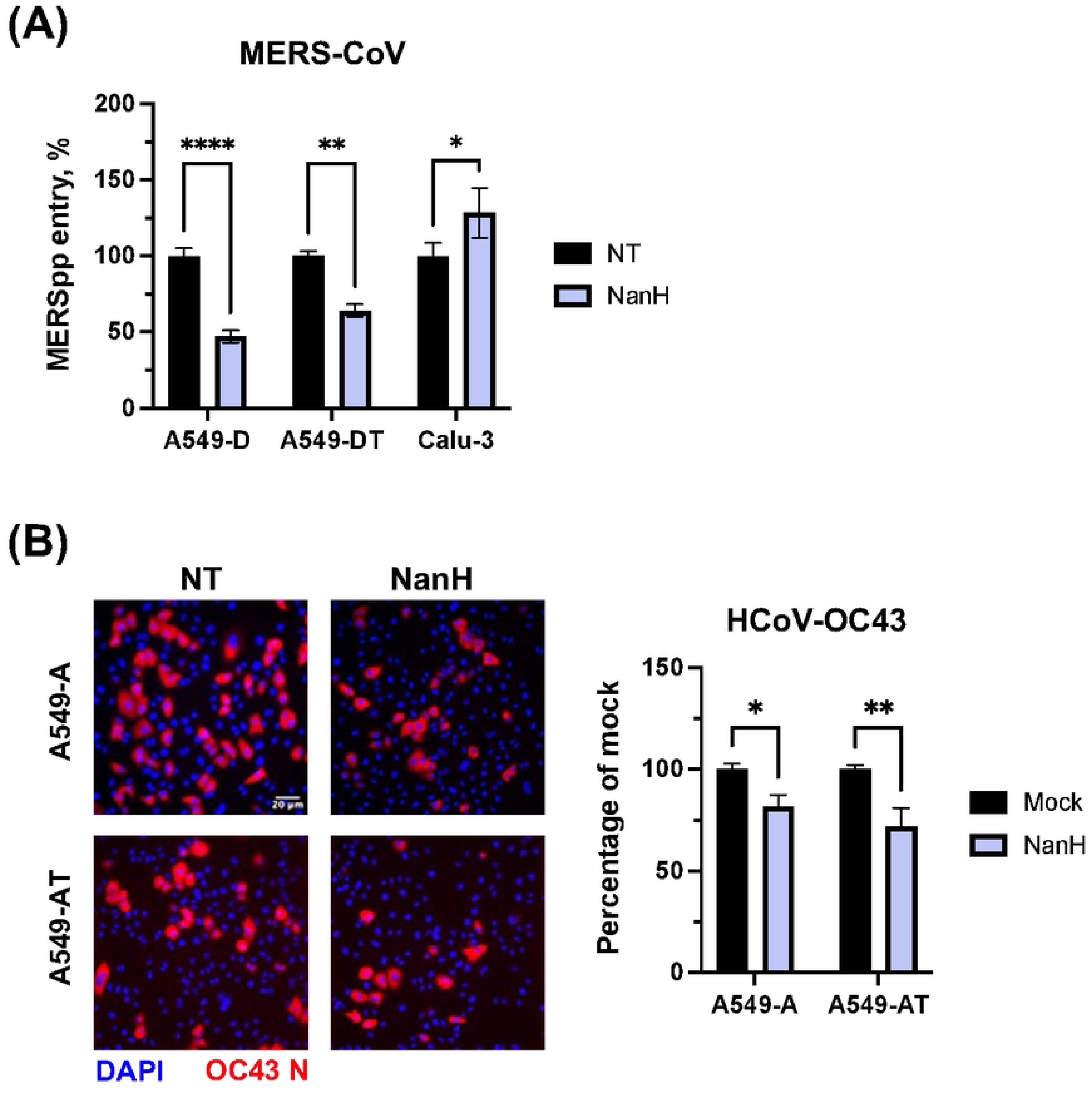
Entry of MERS-CoVpp and HCoV-OC43 is inhibited by the absence of sialic acid regardless of TMPRSS2 expression in A549-derived cells. **(A)** A549-D, A549-DT or Calu-3 cells were pre-treated with NanH diluted to 50 μg/mL in serum-free media for 30 minutes at 37°C. Cells were then washed and inoculated with MERS-CoVpp for 2 h at 37°C prior to being washed and overlaid with complete media for 72 h. Luciferase activity was then measured to assess viral entry. Data are expressed as percentage relative to DMSO control. **(B-C)** A549-A or A549-AT cells were pre-treated with NanH as above **(A)**, then washed and infected with HCoV-OC43 (MOI 0.5) for 1 h at 37°C. Cells were then incubated in complete media for an additional 7 h, at which point they were fixed and processed for immunofluorescence microscopy using an antibody against the HCoV-OC43 N protein. Representative images are shown (20X magnification; scale bar, 20 μm). **(C)** The percentage of infected cells in each condition was determined using ImageJ. **(A, C)** Graphs show mean +/- SEM **(A)** or SD **(C)** from three independent experiments performed in triplicate. Statistical significance was assessed by two-way ANOVA (*p<0.05; **p<0.01; ****p<0.0001).

As a control, we tested entry of authentic HCoV-OC43, which like MERS-CoV is documented to bind sialic acid (39, 61). HCoV-OC43 entry was similarly inhibited by the removal of sialic acid from the surfaces of A549-A and A549-AT cells **(Fig 6B-C)**. However, it is worth noting that HCoV-OC43 entry was inhibited by E64d, but not camostat, treatment in both cell lines **(S4 Fig)**, reflecting a dependence on the endosomal entry route regardless of TMPRSS2 expression. Thus, we cannot rule out in this assay that sialic acid may promote HCoV-OC43 internalization and endocytic entry in addition to cell attachment. Nonetheless, HCoV-OC43 and MERS-CoV entry was similarly reduced in the absence of sialic acid, regardless of TMPRSS2 expression. These findings could support a more direct role for sialic acid in spike binding (33) during MERS-CoV and HCoV-OC43 entry, since TMPRSS2 expression and the availability of the cell surface entry route did not affect their dependency on sialic acid during entry in A549-derived cells.

## Discussion

In this study, we evaluated whether dependence on sialoglycans during CoV cell entry is affected by entry route, which can occur at the cell surface in a TMPRSS2-dependent manner, or in endosomes in a cathepsin-dependent manner. We confirm that entry of SARS-CoV-1 depends on the endosomal route (62) and show that entry of related pre-emergent bat CoVs WIV1-CoV and WIV16-CoV similarly occurs primarily via the endosomal pathway. While WIV1-CoV and WIV16-CoV have been previously established to use ACE2, as we confirm here **(S1 Fig)**, other aspects of their entry mechanisms had not yet been investigated. Their requirement for endosomal entry shown here is consistent with the lack of furin cleavage site in the spike protein of SARS-CoV-1, WIV1-CoV and WIV16-CoV, which precludes pre-activation of spike in the producer cell and likely therefore limits usage of the TMPRSS2-mediated cell surface pathway for entry. Consistent with previous literature, we show that SARS-CoV-2 Hu1 (63) and delta B1.617.2 variant (54), as well as MERS-CoV (64), favour cell surface entry in the presence of TMPRSS2 **(Fig 2B-C)**, while the SARS-CoV-2 omicron B1.1.592 variant (54) demonstrates a preference for endosomal entry **(Fig 2B)**. SARS-CoV-2 Hu1, delta and MERS-CoV were flexible in entry pathway usage **(Fig 2B-C)**, which has been proposed to aid in evasion of antiviral restriction factors (63, 65–67). Importantly, the flexibility in entry route usage by SARS-CoV-2 Hu1, delta and MERS-CoV enabled us to specifically assess the requirement for sialylated glycans during cell surface or endosomal entry in cell lines either expressing or lacking TMPRSS2, respectively. Using both genetic and enzymatic approaches, we compared the effect of sialic acid removal on CoV cell entry in A549-A cells (lacking TMPRSS2) versus A549-AT cells (ectopically expressing TMPRSS2). Ectopic expression of TMPRSS2 in A549-AT cells rescued entry of SARS-CoV-2 Hu1 and delta variant from the inhibitory effect of sialic acid removal **(Fig 5B-C)**.

In contrast, entry of SARS-CoV-2 omicron B.1.1.529, which relies more on endosomal entry even in the presence of TMPRSS2 (54) **(Fig 2B)**, was similarly restricted by the absence of sialic acid regardless of TMPRSS2 expression **(Fig 5D)**. Consistently, the absence of sialic acid in CMAS KO cells profoundly inhibited entry of cathepsin-dependent CoVs (SARS-CoV-2 omicron, SARS-CoV-1, WIV1-CoV and WIV16-CoV) to a similar extent as inhibiting cathepsin L via E64d treatment. Based on these data, we propose a model where sialic acid plays a more significant role during endosomal entry compared to cell-surface entry of SARS-CoV-2 Hu1 and delta **(Fig 7)**.

**Figure 7.**
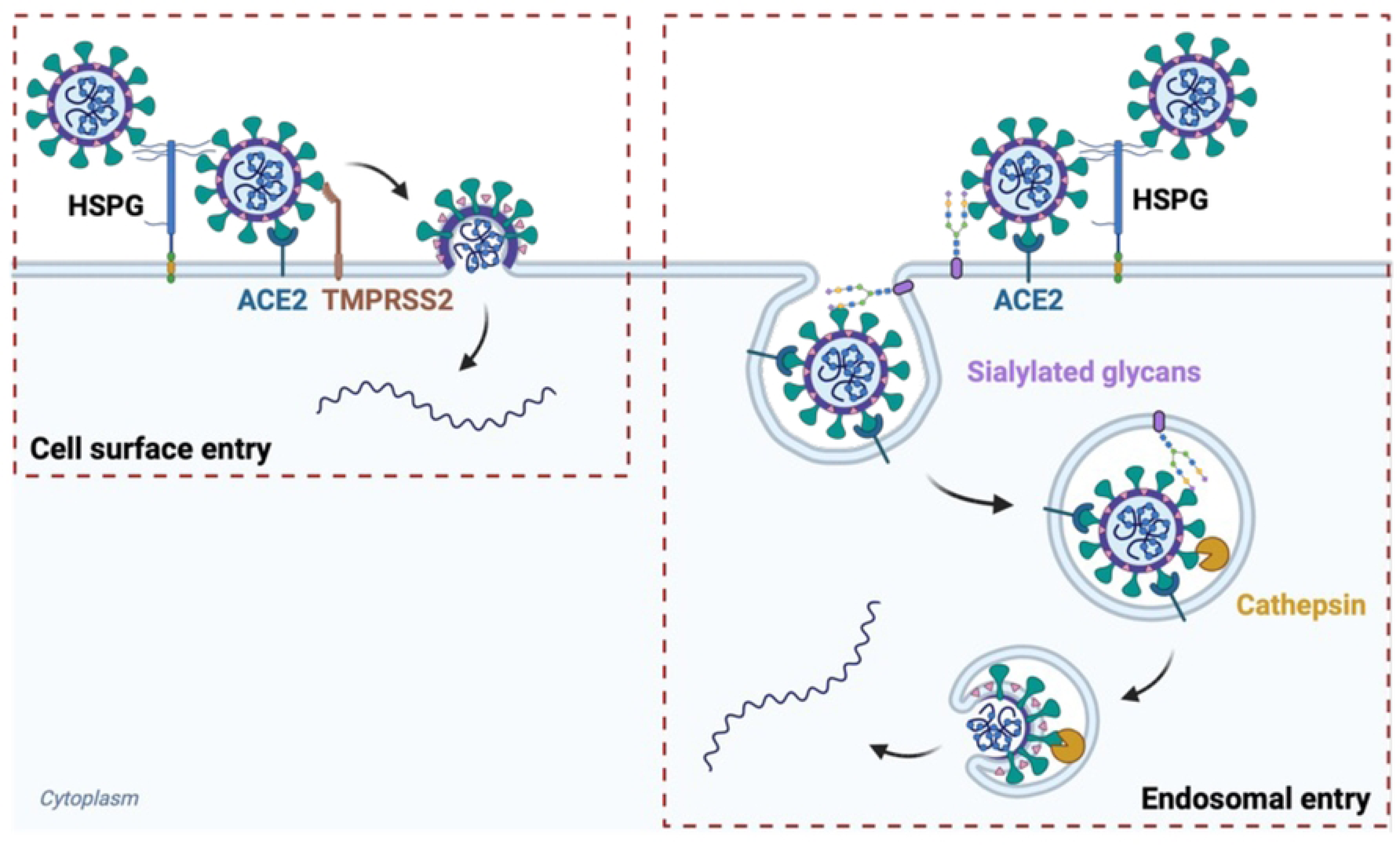
Proposed working model for the role of sialoglycans in SARS-CoV-2 entry. Sialoglycans enhance endosomal cathepsin-mediated entry of SARS-CoV-2, but are not required for cell surface TMPRSS2-dependent entry.

Classically, sialic acid serves as a receptor or co-receptor for some viruses, most notably influenza viruses (60), aiding in attachment of viral particles to cell surfaces. Indeed, the spike proteins of human common cold embecoviruses HCoV-OC43 and HCoV-HKU1 interact with 9-*O*-acetylated sialic acid through a well-conserved receptor binding site in S1, which is required for attachment and entry (28, 31, 68), as we observed for HCoV-OC43 **(Fig 6B-C)**. For HCoV-HKU1, sialoglycan binding promotes the “up” position of S1 (69), which is proposed to enable subsequent entry steps such as binding to a secondary receptor. HCoV-OC43 and HCoV-HKU1 encode hemagglutinin esterase (HE) (70), which serves as a “receptor-destroying enzyme” required for productive infection and release of infectious HCoV-OC43 and HCoV-HKU1 progeny virions (71, 72), similar to other viruses that attach to sialic acid, such as IAV (73), which encodes both receptor binding (hemagglutinin) and receptor-destroying (neuraminidase) enzymatic activities. In contrast, other CoVs, including sarbecoviruses and merbecoviruses, have not evolved to encode HE (70), supporting the notion that entry of these viruses to cells may not rely heavily on sialic acid binding.

We found that binding of SARS-CoV-2 Hu1 spike protein to A549-A cells still occurred in CMAS KO cells or NanH-treated cells lacking sialic acid **(Fig 3B)**, although spike binding is inhibited by heparin treatment **(Fig 3C)**, reflecting the requirement for heparan sulfate proteoglycans in SARS-CoV-2 attachment (26). Consistent with our findings, Hao et al. used glycan microarrays to show that recombinant spike proteins of SARS-CoV-1, SARS-CoV-2 and MERS CoV bind to heparan sulfate, but not to sialylated N-glycans, O-glycans or glycolipid glycans (74). Indeed, MERS-CoV spike protein binding to sialylated glycans was only detected when the S1 domain of spike was displayed on nanoparticles to enable multivalent interactions with increased avidity (33). Even under these conditions, binding of MERS-CoV S1 to sialoglycans occurred with much lower affinity than binding of IAV hemagglutinin (HA) (33), which is well-established to bind α2,6-linked sialic acid (60). Baker et al. also demonstrated binding of recombinant SARS-CoV-2 S1 to multivalent gold nanoparticles bearing sialic acid, which enabled detection of SARS-CoV-2 in a lateral flow assay (75). However, these relatively weak low avidity spike-sialic acid interactions (33, 74, 75) may not contribute greatly to CoV binding to cell surfaces during authentic infection.

Nguyen et al., in contrast, demonstrated that the SARS-CoV-2 Hu1 spike receptor binding domain (RBD) binds to sialic-acid containing glycolipids on ACE2-expressing HEK293 cells (29), which may reflect cell type differences or that exposure of the RBD is required for binding sialylated glycolipids, which is not necessarily recapitulated in our assay with spike SGP. Similarly, Negi et al. used in vitro binding assays to demonstrate binding of SARS-CoV-2 to sialylated gangliosides embedded in supported lipid bilayers (76). As such, spike binding to cell surfaces may be aided by sialic acid in some circumstances. Consistently, compounds containing α2,6-linked sialic acid were shown to inhibit SARS-CoV-2 attachment and infection (36).

Our findings support a functional role for sialic acid in entry of SARS-CoV-2 and MERS-CoV, consistent with prior literature demonstrating that entry and infection of authentic SARS-CoV-2 (29) and MERS-CoV (33) was inhibited in sialidase-treated cells. TMPRSS2 expression in A549 cells enabled SARS-CoV-2, but not MERS-CoV, to overcome the inhibitory effect of sialic acid removal on pseudoparticle entry **(Fig 5B-C)**. However, removal of sialic acid from Calu-3 cells by NanH did not affect the entry of MERS-CoV pseudoparticles, which contrasts a previous study showing that entry of authentic MERS-CoV was inhibited in sialidase-treated Calu-3 cells, but not Vero cells, which the authors attributed to differences in glycan expression levels between the cell types (33). Indeed, differences in protein receptor abundance, protease expression, or glycosylation levels in different cell types or cells cultured in different labs may impact the dependency on sialic acid during MERS-CoV entry. The discrepancy in these findings may also reflect differences between lentiviral pseudoparticles and authentic virus, where differences in the egress route and presence of the envelope and membrane protein during authentic viral infection could affect proteolytic processing, thus affecting entry route preference. While we were unable to perform studies with authentic SARS-CoV-2 or MERS-CoV isolates due the lack of BSL3 facilities, future studies should aim to address the role of sialoglycans during cell-surface or endosomal entry of authentic SARS-CoV-2 and MERS-CoV isolates.

We propose a model where sialic acid plays a role in promoting the internalization of SARS-CoV-2 when the endocytic entry pathway is used. Prior literature supports roles for sialic acid in post-binding viral entry steps. For example, using Lec1 cells, a mutant CHO cell line that is unable to synthesize complex and hybrid N-linked glycans and thus lacks sialylated-N-glycans, Chu & Whittaker demonstrated that terminal N-linked glycosylation was required for influenza virus internalization, but not binding or fusion (37). Similarly, adeno-associated virus still binds to CHO cells lacking surface expression of sialic acid, but it does not undergo internalization (77). In Huh7 hepatoma cells, Zika virus internalization, but not attachment, requires α2,3-linked sialic acid (38). Huh7 cells deficient in α2,3-linked sialic acid were also less susceptible to MERS-CoV pseudoparticle infection (38), which could also indicate a role for sialic acid in MERS-CoV internalization as we found that the endosomal entry pathway is preferred during MERS-CoV entry in Huh7 cells (data not shown). Interestingly, we observed that entry of VSVpp, like that of CoVpp, was inhibited in the CMAS KO cells, which could reflect a proposed role for sialic acid in VSV attachment (59) or a general defect in endocytosis, which is also required for VSV fusion and entry (78). The latter would be consistent with literature showing that endocytosis is reduced in dendritic cells upon removal of terminal α2,3 or α2,6-linked sialic acid from the cell surface (79). However, future studies are necessary to examine the specific impact of sialic acid removal on endocytosis in epithelial cell lines and to investigate the underlying mechanisms.

In conclusion, our study has contributed to understanding the roles of sialoglycans in CoV entry, suggesting a role for sialic acid in aiding endosomal entry of SARS-CoV-2. While direct binding of sialic acid by spike may contribute to entry of some CoVs, such as embecoviruses and merbecoviruses, additional roles of sialic acid in promoting virion internalization may enhance entry through the endosomal route. Given that clinical CoV isolates are often more dependent on cell surface TMPRSS2-mediated entry, antiviral efficacy of sialic acid-derived inhibitors should be evaluated in the context of both entry routes. Overall, these findings contribute to understanding CoV entry and may inform design of antiviral molecules that act by disrupting CoV-glycan interactions.

## Materials and Methods

### Plasmids

CoV lentiviral pseudoparticles (CoVpp) were produced using packaging, transfer, and CoV spike plasmids. The packaging plasmids used include pHDM-Hgpm2 encoding HIV-1 gag-pol (BEI Resources NR-52517), pHDM-tat1b encoding HIV-1 tat (BEI Resources #NR-52518), and pRC-CMV-rev1b encoding HIV-1 rev (BEI Resources #NR-52517). Alternatively, psPAX-2 (Addgene #12260) was used as a packaging plasmid. Transfer plasmids pHAGE-Luc-ZsGreen (BEI Resources #NR-52519) or pLenti-CMV-Puro-Luc (Addgene #17477) were used. SARS-CoV-2 spike expression plasmids that were used include pHDM encoding full length (FL) SARS-CoV-2 Wuhan-Hu-1 spike (BEI Resources, #NR-52514) and pcDNA3.1 encoding a 19 amino acid C-terminally truncated (ΔCT) SARS-CoV-2 Wuhan Hu-1 spike obtained from Dr. Raffaele De Francesco (Addgene #155297). For delta and omicron variants, plasmids pTwist-SARS-CoV-2 d18 B.1.617.2v1 (Addgene #179905) and pTwist-SARS-CoV-2 d18 B.1.1.529 (Addgene #179907) were kindly shared by Dr. Alejandro Balazs. We previously synthesized pcDNA3.1-SARS1-S encoding SARS-CoV-1 spike protein (Tor2 strain, GenBank accession no. <u>NC_004718.3</u>) and pcDNA3.1-WIV1-S encoding WIV1-CoV spike protein (GenBank accession no. <u>KC881007.1</u>) (80). Plasmids pCAGGS-MERS-S (81) and pEVAC-WIV16-S (82) plasmids, encoding MERS-CoV and WIV16-CoV spike proteins, respectively, were also described previously.

For generation of stable cell lines via lentiviral transduction, the packaging plasmid psPAX2 was used along with pCMV-VSV-G encoding the VSV G glycoprotein (Addgene #8454). Depending on the desired modification, different transfer plasmids were used. For the generation of the A549-ACE2 and A549-DPP4 cell lines, lentiviral vectors encoding human ACE2 (Dr. Sonja Best, Addgene #154981) and DPP4 (Addgene #158452) was used, respectively. TMPRSS2 was subcloned into the pLJM1 lentiviral vector (Addgene #19319) by standard cloning approaches, with pLJM1-GFP used as a control.

For mammalian protein expression of soluble spike glycoprotein and the CR3022 anti-spike antibody, the following reagents were obtained through BEI Resources: Modified pαH vector containing the SARS-CoV-2 Wuhan-Hu-1 HexaPro spike glycoprotein ectodomain (#NR-53587), and the plasmid set for Anti-SARS Coronavirus Human Monoclonal Antibody CR3022 (#NR-53260).

### Cells and viruses

HEK293T/17 (ATCC #ACS-4500) and HCT-8 (ATCC #CCL-244) cells were cultured in Dulbecco’s modified Eagle medium (DMEM; ThermoFisher #11995065) supplemented with 10% FBS, 50 U/mL penicillin, and 50 µg/mL streptomycin. ExpiCHO cells (ThermoFisher) were cultured in ExpiCHO Expression Medium (ThermoFisher) and were maintained in a humid 5% CO_2_ atmosphere at 37 °C with shaking at 120 rpm. Calu-3 (ATCC HTB-55) cells were cultured in minimal essential medium (MEM; Fisher Scientific #11095080) supplemented with 10% FBS, 1% sodium pyruvate, 1% non-essential amino acid solution, 50 U/mL penicillin, and 50 µg/mL streptomycin. Parental A549 cells (BEI Resources #NR-52268) were cultured in Ham’s F-12 K (Kaighn’s) medium (Fisher Scientific #21127030) with 10% FBS, 50 U/mL penicillin, and 50 µg/mL streptomycin. A549-ACE2 and A549-DPP4 cells were cultured in complete Ham’s F-12K media supplemented with 10 µg/mL blasticidin. A549-ACE2-GFP and A549-ACE2-TMPRSS2 cells were cultured in Ham’s F-12 K (Kaighn’s) medium supplemented with 10% FBS, 1% pen/strep, 10 µg/mL blasticidin and 2 µg/mL puromycin. All cell lines were maintained at 37 °C and 5% CO_2._

A549-ACE2 and A549-DPP4 cells were generated by lentiviral transduction and selected for in normal A549 media supplemented with 10 µg/mL of blasticidin. The bulk population of A549-ACE2 cells was single-cell cloned as described previously (80) and ACE2 expression was re-confirmed by western blot. DPP4 expression in the bulk population was similarly confirmed by western blot.

HCoV-OC43 (BEI Resources NR-52725) was propagated in HCT-8 cells as we have described previously (80). Influenza A virus (IAV) (strain A/New York/18/2009, H1N1) was obtained from BEI Resources (NR-15268) and propagated in Madin-Darby canine kidney cells (BEI Resources NR2628) in the lab of Dr. Katrina Gee (Queen’s University).

### Antibodies

For western blot, we used a rabbit monoclonal antibody against ACE2 (ThermoFisher #MA5-32307; diluted 1:1000), a rabbit monoclonal antibody against DPP4 (ThermoFisher #MA5-32643; diluted 1:1000) and a mouse monoclonal antibody against GAPDH as a loading control (ThermoFisher #MA5-15738). Secondary antibodies for western blot were goat anti-rabbit IgG IRDye 800CW Conjugate (LI-COR #926-32211) or goat anti-mouse IgG IRDye 680RD conjugate (LI-COR #926-68070) and membranes visualized on a LICOR Odyssey CLx as described (80). For immunofluorescence, a primary antibody against the HCoV-OC43 N protein (Millipore #MAB9013; diluted 1:500) was used, followed by a secondary Alexa Fluor 555 conjugated anti-mouse IgG antibody (NEB #4413S; diluted 1:1000). For flow cytometry, the following primary antibodies were used: Phycoerythrin (PE)-conjugated anti-human TMPRSS2 antibody (BioLegend #378403) and CR3022 (generated as described below). For the latter, a secondary Alexa Fluor 647 conjugated goat anti-human IgG (H+L) antibody (Invitrogen #A-21445; 10 μg/mL) was used.

### Inhibitors

The TMPRSS2 inhibitor camostat mesylate was obtained from Sigma Aldrich (SML0057) and the cathepsin L inhibitor E64d was obtained from Tocris Bioscience (Cat# 4545). Stocks were prepared in DMSO, which was used as a vehicle control.

### Enzyme expression and purification

The gene sequence of *C. perfringens* α2-3,-6,-8 neuraminidase (NanH) (GenBank: Y00963.1) was commercially synthesized and ligated into a pET-15b plasmid using NdeI and XhoI restriction sites (Genscript). The resulting vectors were chemically transformed into *Escherichia coli* BL-21 competent cells. Transformed cells were cultured in 5 mL LB media supplemented with 100 μg/mL ampicillin for 16 h at 37 ℃ with shaking (200 rpm). Cultured cells were then scaled up to 500 mL in LB-amp to an optical density (OD_600_) of 0.8. Protein expression was then induced with addition of isopropyl β-D-1-thiogalactopyranoside (IPTG) to a final concentration of 0.1 mM and cells were incubated at 25 ℃ with shaking (200 rpm) for 16 hours before harvesting by centrifugation. Pelleted cells were resuspended in lysis buffer (0.1 M Tris-HCl, pH 8.0, 0.1% Triton X-100) and lysed by high pressure homogenization using an Avestin Emulsiflex C3. Cell debris was removed by ultracentrifugation, and the supernatant was mixed 1:1 with binding buffer (10 mM imidazole, 500 mM NaCl, 50 mM Tris-HCl, pH 7.5). The resulting solutions were then filtered using a 0.22 μm filter unit and applied to a Ni^2+^-sepharose resin (GE Healthcare, fast flow) pre-equilibrated with binding buffer. The column was washed with 6 column volumes (CV) of binding buffer, 6 CV wash buffer (50 mM imidazole, 500 mM NaCl, 50 mM Tris–HCl, pH 7.5), and eluted with 10 CV elution buffer (200 mM imidazole, 500 mM NaCl, 50 mM Tris-HCl, pH 7.5). Overexpression and purity were assessed by SDS-PAGE. Purified NanH was buffer exchanged into 20 mM Tris-HCl (pH 7.5) buffer and concentrated using a 10 kDa MWCO spin filter (Amicon). NanH was stored in 20 mM Tris-HCl (pH 7.5) with 20% glycerol at −80°C. Final protein concentrations were determined by BCA assay (ThermoFisher), and the yield of NanH was 175 mg/500 mL culture.

### Recombinant protein expression and purification

For expression of soluble spike glycoprotein or CR3022 anti-SARS antibody, plasmids were first chemically transformed into DH5α competent cells. Cells containing each plasmid were cultured in 5 mL LB media supplemented with appropriate antibiotics (100 μg/mL ampicillin for soluble spike protein ectodomain, 50 μg/mL zeocin for CR3022 heavy chain, 100 μg/mL blasticidin for CR3022 light chain) for 16 h at 37 °C with shaking (200 rpm) for 16 hours. Cultured cells were then scaled up to 500 mL and plasmids were isolated using the PureLink™ HiPure Plasmid Maxiprep Kit (ThermoFisher #K210006). The isolated plasmids were filter-sterilized prior to transfection. ExpiCHO cells were transiently transfected with the vector containing the gene encoding soluble spike glycoprotein ectodomain or CR3022 (using a heavy chain:light chain ratio of 1:2) using the Expifectamine 293 Transfection Kit (ThermoFisher #A29129), following the user manual. The cells were monitored throughout the expression and the culture was harvested by centrifugation (25 min, 4000 rcf) on day 7 following transfection.

The soluble spike glycoprotein ectodomain was purified using its C-terminal octa-histidine tag. After centrifugation, the supernatant was collected and adjusted to contain 20 mM imidazole, 200 mM NaCl, and 30 mM sodium phosphate at pH 7.2. The resulting solutions were then filtered using a 0.45 μm PES Filter Unit and applied to a Ni^2+^-sepharose resin (Cytiva) pre-equilibrated with column buffer (20 mM HEPES, 300 mM NaCl, pH 7.2) containing 20 mM imidazole. The column was sequentially washed with column buffer containing 20 mM, 50 mM, and 100 mM imidazole prior to elution with 300 mM imidazole. Overexpression and purification were assessed by SDS-PAGE. Purified protein was buffer exchanged into 20 mM Tris-HCl (pH 8.0) with 500 mM NaCl, concentrated using a 30 kDa MWCO spin filter (Amicon), and stored in 20 mM Tris-HCl (pH 8.0) with 500 mM NaCl_2_ at −80 °C in small aliquots. The final protein concentration was determined by BCA assay (ThermoFisher), and the yield was 3.44 mg/100 mL culture.

The CR3022 was purified by protein G affinity column using its inherent human IgG1 Fc region. After centrifugation, the supernatant was collected and adjusted to contain binding buffer (20 mM sodium phosphate at pH 7.0). The resulting solutions were then filtered using a 0.45 μm PES Filter Unit and purified with a HiTrap protein G HP column (Cytiva) according to the manual. The filtered solution was applied to the column pre-equilibrated with binding buffer and sequentially washed with a 10-column volume of binding buffer. To elute bound antibodies, 0.1 M glycine-HCl at pH 2.7 was applied to the column, and fractions were collected in tubes containing 1 M Tris-HCl at pH 9.0 for immediate neutralization. Overexpression and purification were assessed by SDS-PAGE. Purified protein was buffer exchanged into PBS, concentrated using a 30 kDa MWCO spin filter (Amicon), and stored in PBS at −80 °C in small aliquots to avoid repeated freeze-thaw. The final protein concentration was determined by BCA assay (ThermoFisher), and the yield of CR3022 was 1.28 mg/100 mL culture.

Fluorescein-conjugated *Sambucus nigra* lectin (SNA) that binds sialic acid and *Erythrina cristagalli* lectin (ECL) that binds galactose were acquired from Vector Laboratories (SA-5001-1 and FL-1141-5, respectively).

### Pseudoparticle entry assays

Lentiviral pseudoparticles were generated as described previously (36) with minor modifications. Using Lipofectamine 2000 (Thermo Fisher Scientific 11668030) HEK-293T/17 cells were co-transfected with packaging, transfer, and spike plasmids. Cell culture supernatants containing the pseudoparticles was collected at 48 and 72 h post-transfection, pooled, filtered using a 0.45 µm filter, and aliquoted for storage at −80°C.

CoV pseudoparticles (CoVpp) were used to infect target cells seeded in triplicate in 96-well plates. Cells were inoculated with pseudoparticles for 2 h and then replaced with the appropriate media. Infectivity was determined after 3 days by luminescence using a Promega GloMax plate reader following addition of BrightGlo reagent (Promega #E2620) to Calu-3 cells or FLuc Assay buffer (Nanolight #318) supplemented with 0.25% Triton-X100 to all other cells. For inhibitor experiments, cells were pre-treated with 25 µM camostat mesylate or 10 µM E64d for 1 h at 37°C prior to inoculation with CoVpp.

### HCoV-OC43 entry assays

A549-ACE2-GFP or A549-ACE2-TMPRSS2 cells were plated at a density of 10,000 cells/well in 96-well plates. The following day, cells were treated with diluted NanH or protease inhibitors, then washed and infected with HCoV-OC43 (MOI 0.5) for 1 h at 37°C. The inocula were removed, and cells were washed and incubated in complete media for an additional 7 h. Cells were then fixed using 10% formalin and processed for immunofluorescence using an antibody against the HCoV-OC43 N protein (Millipore #MAB9013; diluted 1:500) and a secondary Alexa Fluor 555 conjugated anti-mouse IgG antibody (NEB #4413S; diluted 1:1000). The percentage of infected cells was calculated using an open-source Fiji macro (83).

### Enzyme pre-treatment assays

For enzyme pre-treatment assays, NanH enzyme stocks were diluted to 50 μg/mL in serum-free DMEM. Cells were pretreated with NanH for 30 minutes at 37°C prior to staining with FITC-conjugated lectins (SNA-FITC or ECL-FITC), or inoculation with CoVpp, or IAV. Lectin staining was visualized via fluorescence microscopy using a Nikon Eclipse Ts2-FL inverted microscope. Images were captured using a Nikon DS-Fi3 6MP camera. CoVpp entry was assessed by luciferase reporter activity after 72 h. Infection of Calu-3 cells at 3×10^5^, 3×10^4^, or 3×10^3^ plaque forming units (pfu) of IAV was assessed by measuring IAV M RNA expression via RT-qPCR after 8 h.

### Flow cytometry

Adherent cells were first washed three times with PBS without Ca^2+^/Mg^2+^ and detached using cell dissociation buffer (Gibco #13151014) for 10 min at 37 °C. Once cells were visibly detached, serum-free media was added, and the cells were gently centrifuged (300 rpm for 3 min) and resuspended to have 0.25 million cells per sample in FACS buffer (PBS without Ca^2+^/Mg^2+^ supplemented with 2 mM EDTA and 0.5% BSA for TMPRSS2 expression and SGP binding experiments and 1 % FBS/PBS for lectin binding experiments). Cells were centrifuged gently and washed twice with respective buffers.

For the detection of cell-surface TMPRSS2 expression, washed cells were resuspended in 100 μL of FACS buffer with 5 μL of PE anti-human TMPRSS2 antibody (BioLegend #378403) per million cells and incubated for 30 min at 4 °C in the dark. For the detection of cell-surface sialic acid and confirmation of sialic acid cleavage by NanH, washed cells were resuspended in 100 uL of 1% FBS/PBS with 20 μg/mL SNA-FITC or 5 μg/mL ECL-FITC for 30 min at 4 °C in the dark. For the detection of SGP binding, washed cells were resuspended in 100 μL of FACS buffer with 5 μg/mL SGP and incubated for 30 min at 4 °C. For heparin treatment, SGP was incubated with 10 μg/mL heparin in FACS buffer for 10 min prior to its addition to cells. CR3022 antibody (10 μg/mL) and 10 μg/mL of goat anti-human IgG (H+L) secondary antibody Alexa Fluor 647 (Invitrogen #A-21445) was pre-incubated for 30 min at 4 °C. After 30 min, cells with SGP were centrifuged gently, washed twice with FACS buffer and incubated with 100 uL of the pre-complexed antibodies for 30 min at 4 °C in the dark.

For the TMPRSS2 expression and SGP binding experiments, stained cells were then washed, stained with fixable viability stain, and fixed (Invitrogen #L23101) following the user manual. After fixation, the cells were centrifuged gently, and resuspended in 300 μL of FACS buffer and transferred to U-bottom 96-well plates for flow cytometric analysis (Beckman Coulter, Cytoflex S). The live population of cells was gated based on forward and side scatter emission and exclusion of viability stain positive cells on the FITC (525/40 BP filter) emission channel. Anti-TMPRSS2 or SGP binding were determined by fluorescence intensity on PE (660/20 BP filter) or APC (660/20 BP) emission channel, respectively.

For the lectin binding experiments, stained cells were washed/centrifuged three times with 1 % FBS/PBS, centrifuged gently, and resuspended in 300 μL of FACS buffer and transferred to U-bottom 96-well plates for flow cytometric analysis (Beckman Coulter, Cytoflex S). Cell viability was determined by adding 1 μg/mL propidium iodide (PI) to cell suspensions 1 min prior to analysis. The live population of cells was gated based on forward and side scatter emission and exclusion of PI positive cells on the ECD (610/20 BP filter) emission channel. Lectin binding was determined by fluorescence intensity on the FITC (525/40 BP filter) emission channel.

### Statistical Analysis

Data are represented as mean ± standard deviation or standard error of the mean. Statistical significance was assessed using unpaired t-test with Welch’s correction, or using one-way or two-way ANOVA using Prism 9 (GraphPad Software).

## Acknowledgements

This work was supported by Discovery Grants from the Natural Sciences and Engineering Research Council of Canada (CCC, CJC), the J.P. Bickell Foundation for Medical Research (CCC, CJC), the Canadian Institutes for Health Research (CCC) and a COVID-19 rapid response grant from Queen’s University (CJC, CCC). We are grateful to the Canadian Foundation for Innovation John R. Evans Leaders Fund (CCC) for equipment supporting this project. EVL was supported by a Vanier Canada Graduate Scholarship (EVL), while YK acknowledges an NSERC postgraduate scholarship. We are grateful to Emma Kalin and Dr. Katrina Gee (Queen’s University) for providing influenza A virus.

## Supplementary Figure Captions

**Supp. Figure 1. ACE2 dependence of WIV1-CoV and WIV16-CoV. (A)** Western blot assessing ACE2 expression in parental A549 cells and A549-ACE2 cells. **(B)** A549 or A549-ACE2 cells were inoculated with lentiviral particles pseudotyped with the spike proteins of WIV1-CoV or WIV16-CoV for 2 h, then incubated for an additional 72 h, at which point luciferase activity was measured to assess pseudoparticle entry. The data are expressed as fold change relative to the luciferase signal obtained with no envelope. Graphs show mean +/- SEM from three independent experiments performed in triplicate.

**Supp. Figure 2. Entry of full-length spike pseudotyped lentiviral particles is enhanced in Calu-3 cells related to C-terminally truncated spike pseudotyped particles.** Calu-3 cells were inoculated with lentiviral particles pseudotyped with the spike proteins of SARS-CoV-2 Hu1 (full-length or C-terminally truncated) or pseudoparticles lacking envelope protein (no env) for 2 h, then incubated for an additional 72 h, at which point luciferase activity was measured to assess pseudoparticle entry. The data are expressed as fold change relative to the luciferase signal obtained with no envelope. Graphs show mean +/- SEM from three independent experiments performed in triplicate.

**Supp. Figure 3. NanH effectively removes sialic acid from A549 and Calu-3 cells. (A-B)** A549 cells or Calu-3 cells were pre-treated with NanH diluted to 50 μg/mL in serum-free media for 30 minutes at 37°C. NanH-treated cells were stained with SNA-FITC (binds sialic acid) or ECL-FITC (binds galactose) diluted to final concentration of 20 μg/mL in PBS, then washed with PBS and imaged by fluorescence microscopy (representative images shown in Figure 5A). The mean FITC fluorescence intensity was calculated using ImageJ and is plotted in the graphs. **(A)** NanH-treated Calu-3 cells were infected with the indicated doses of IAV. At 8 hpi, cellular lysates were collected and IAV RNA (encoding the M gene) was assessed by RT-qPCR. The data were normalized to actin and are expressed relative to the non-treated condition. Graphs show mean +/- SD (A) or SEM **(B)** from three independent experiments performed in triplicate. Statistical significance was assessed by two-way ANOVA (***p<0.001; ****p<0.0001).

**Supp. Figure 4. HCoV-OC43 entry requires endosomal cathepsins, but not TMPRSS2. (A-B)** A549-A or A549-AT cells were pre-treated for 1 h at 37°C with DMSO, camostat (25 μM) or E64d (10 μM) diluted in media, then infected with HCoV-OC43 at an MOI of 0.5 ffu/cell for 1 h at 37°C. Cells were then incubated in complete media for an additional 7 h, at which point they were fixed and processed for immunofluorescence microscopy using an antibody against the HCoV-OC43 N protein. **(A)** Representative images are shown (20X magnification; scale bar, 20 μm). **(B)** The percentage of infected cells in each condition was determined using ImageJ. Graphs show mean +/- SEM from three independent experiments performed in triplicate. Statistical significance was assessed by two-way ANOVA (ns, not significant; ***p<0.001; ****p<0.0001).

